## supplementary data for "Divergence in a Eukaryotic Transcription Factor’s co-TF Dependence Involves Multiple Intrinsically Disordered Regions"

**Supplementary Table 1. Plasmids used in this study**

| ID | Plasmid | Derived | Source | Comment |
|---|---|---|---|---|
| pH246 | pGAD-C3 | | PMID:8978031, gift from Fassler lab | Yeast two-hybrid |
| pH247 | pGBD-C3 | | PMID:8978031, gift from Fassler lab | Yeast one-hybrid |
| pH343 | pRS306-GAL1pr-mCherry | pRS306 | this study | Yeast one-hybrid |
| pH346 | pRS314-GAL4pr-Gal4 | pRS314 | this study | Yeast one-hybrid |
| pH349 | pGBD-Gal4DBD-ScE1 | pH247 | this study | Yeast one-hybrid |
| pH350 | pGBD-Gal4DBD-CgE1 | pH247 | this study | Yeast one-hybrid |
| pH351 | pGBD-Gal4DBD-ScE1-ScAD | pH247 | this study | Yeast one-hybrid |
| pH352 | pGBD-Gal4DBD-CgE1-CgAD | pH247 | this study | Yeast one-hybrid |
| pH362 | pGBD-Gal4DBD-CgAD | pH247 | this study | Yeast one-hybrid |
| pH363 | pGBD-Gal4DBD-ScAD | pH247 | this study | Yeast one-hybrid |
| pH377 | pGBD-Gal4DBD-CgE2 | pH247 | this study | Yeast one-hybrid |
| pH378 | pGBD-Gal4DBD-ScE2 | pH247 | this study | Yeast one-hybrid |
| pH379 | pGBD-Gal4DBD-ScAD.9aa | pH247 | this study | Yeast one-hybrid |
| pH380 | pGBD-Gal4DBD-CgAD.2 | pH247 | this study | Yeast one-hybrid |
| pH386 | pGBD-Gal4DBD-CgE2.9aa | pH247 | this study | Yeast one-hybrid |
| pH387 | pGBD-Gal4DBD-CgAD.1 | pH247 | this study | Yeast one-hybrid |
| pH388 | pGBD-Gal4DBD-CgE1-ScAD.9aa | pH247 | this study | Yeast one-hybrid |
| pH389 | pGBD-Gal4DBD-ScAD.9aa-CgE1 | pH247 | this study | Yeast one-hybrid |
| pH390 | pGBD-Gal4DBD-ScAD.9aa-CgE2.9aa | pH247 | this study | Yeast one-hybrid |
| pH391 | pGBD-Gal4DBD-CgE2.9aa-ScAD.9aa | pH247 | this study | Yeast one-hybrid |
| pH434 | pGBD-Gal4DBD-ScPho2AD | pH247 | this study | Yeast one-hybrid |
| pH394 | pGAD-ScPho4ΔDBD | pH246 | this study | Yeast two-hybrid |
| pH395 | pGBD-Pho2Pho4int | pH247 | this study | Yeast two-hybrid |
| pH396 | pGAD-CgPho4ΔDBD | pH246 | this study | Yeast two-hybrid |
| pH050 | pET-11a-ScPho4 DBD-6xHis | pET11a | this study | BLI |
| pH051 | pET-11a-CgPho4 DBD-6xHis | pET11a | this study | BLI |
| bH404 | pET-11a-GST-CgPho4 | pET11a | this study | PBM |
| pH073 | bRA89, Cas9, CEN/ARS, HygR | | PMID: 28405019 gift from Malkova lab | CRISPR |
| pH173 | ScPHO4pr, mNeon, CEN/ARS, LEU2 | pRS315 | this study | Chimera backbone |
| pH188 | Cg(1-533) | pH173 | this study | Chimera |
| pH194 | Sc(1-312) | pH173 | this study | Chimera |

| | | | | |
|---|---|---|---|---|
| pH209 | Cg(1-112) Sc(100-176) Cg(283-533) | pH173 | this study | Chimera |
| pH210 | Cg(1-458) Sc(243-312) | pH173 | this study | Chimera |
| pH211 | Cg(1-282) Sc(177-242) Cg(459-533) | pH173 | this study | Chimera |
| pH212 | Sc(1-42) Cg(45-533) | pH173 | this study | Chimera |
| pH213 | Cg(1-44) Sc(43-99) Cg(113-533) | pH173 | this study | Chimera |
| pH215 | Sc(1-99) Cg(113-533) | pH173 | this study | Chimera |
| pH216 | Sc(1-42) Cg(45-458) Sc(243-312) | pH173 | this study | Chimera |
| pH217 | Cg(1-44) Sc(43-176) Cg(283-533) | pH173 | this study | Chimera |
| pH218 | Cg(1-112) Sc(100-242) Cg(459-533) | pH173 | this study | Chimera |
| pH219 | Cg(1-282) Sc(177-312) | pH173 | this study | Chimera |
| pH220 | Sc(1-42) Cg(45-112) Sc(100-176) Cg(283-533) | pH173 | this study | Chimera |
| pH221 | Cg(1-44) Sc(43-99) Cg(113-282) Sc(177-312) | pH173 | this study | Chimera |
| pH222 | Sc(1-42) Cg(45-282) Sc(177-312) | pH173 | this study | Chimera |
| pH223 | Cg(1-44) Sc(43-99) Cg(113-282) Sc(177-242) Cg(459-533) | pH173 | this study | Chimera |
| pH224 | Sc(1-42) Cg(45-112) Sc(100-176) Cg(283-458) Sc(243-312) | pH173 | this study | Chimera |
| pH227 | Sc(1-176) Cg(283-533) | pH173 | this study | Chimera |
| pH229 | Sc(1-176) Cg(283-458) Sc(243-312) | pH173 | this study | Chimera |
| pH230 | Sc(1-99) Cg(113-282) Sc(177-312) | pH173 | this study | Chimera |
| pH231 | Cg(1-44) Sc(43-312) | pH173 | this study | Chimera |
| pH232 | Sc(1-42) Cg(45-112) Sc(100-312) | pH173 | this study | Chimera |
| pH233 | Sc(1-242) Cg(459-533) | pH173 | this study | Chimera |
| pH234 | Cg(1-112) Sc(100-312) | pH173 | this study | Chimera |
| pH235 | Sc(1-99) Cg(113-458) Sc(243-312) | pH173 | this study | Chimera |
| pH236 | Cg(1-76) Sc(71-99) Cg(113-282) Sc(177-312) | pH173 | this study | Chimera |
| pH237 | Cg(1-44) Sc(43-70) Cg(77-282) Sc(177-312) | pH173 | this study | Chimera |
| pH239 | Sc(1-99) Cg(113-282) Sc(177-242) Cg(459-533) | pH173 | this study | Chimera |
| pH240 | Cg(1-112) Sc(100-176) Cg(283-458) Sc(243-312) | pH173 | this study | Chimera |

| | | | | |
|---|---|---|---|---|
| pH241 | Cg(1-44) Sc(43-242) Cg(459-533) | pH173 | this study | Chimera |
| pH250 | Cg(1-44) Sc(43-176) Cg(283-458) Sc(243-312) | pH173 | this study | Chimera |
| pH251 | Sc(1-42) Cg(45-282) Sc(177-242) Cg(459-533) | pH173 | this study | Chimera |
| pH252 | Sc(1-42) Cg(45-112) Sc(100-242) Cg(459-533) | pH173 | this study | Chimera |
| pH253 | Cg(1-44) Sc(43-99) Cg(113-458) Sc(243-312) | pH173 | this study | Chimera |
| pH254 | Cg(1-44) Sc(43-176) Cg(283-327) Sc(205-242) Cg(459-533) | pH173 | this study | Chimera |
| pH255 | Cg(1-44) Sc(43-204) Cg(328-533) | pH173 | this study | Chimera |
| pH256 | Cg(1-44) Sc(43-220) Cg(377-533) | pH173 | this study | Chimera |
| pH257 | Cg(1-282) Sc(177-204) Cg(328-533) | pH173 | this study | Chimera |
| pH258 | Cg(1-327) Sc(205-242) Cg(459-533) | pH173 | this study | Chimera |
| pH265 | Sc(1-204) Cg(328-533) | pH173 | this study | Chimera |
| pH266 | Sc(1-153) Cg(250-533) | pH173 | this study | Chimera |
| pH276 | Sc(1-42) Cg(45-282) Sc(177-204) Cg(328-533) | pH173 | this study | Chimera |
| pH277 | Cg(1-44) Sc(43-99) Cg(113-282) Sc(177-204) Cg(328-533) | pH173 | this study | Chimera |
| pH278 | Sc(1-204) Cg(328-458) Sc(243-312) | pH173 | this study | Chimera |
| pH279 | Cg(1-44) Sc(43-204) Cg(328-458) Sc(243-312) | pH173 | this study | Chimera |
| pH294 | Cg(1-470) Sc(251-312) | pH173 | this study | Chimera |
| pH301 | Sc(1-247) Cg(464-533) | pH173 | this study | Chimera |
| pH326 | Sc(1-156) Cg(253-533) | pH173 | this study | Chimera |
| pH327 | Cg(1-252) Sc(157-312) | pH173 | this study | Chimera |
| pH328 | Sc(1-42) Cg(45-252) Sc(157-312) | pH173 | this study | Chimera |
| pH329 | Cg(1-44) Sc(43-247) Cg(464-533) | pH173 | this study | Chimera |
| pH330 | Cg(1-44) Sc(43-156) Cg(253-463) Sc(251-312) | pH173 | this study | Chimera |
| pH331 | Cg(1-44) Sc(43-156) Cg(253-533) | pH173 | this study | Chimera |
| pH332 | Sc(1-42) Cg(45-463) Sc(251-312) | pH173 | this study | Chimera |
| pH334 | Sc(1-42) Cg(45-252) Sc(157-247) Cg(464-533) | pH173 | this study | Chimera |

| | | | | |
|---|---|---|---|---|
| pH436 | Cg(1-112) Sc(100-176) Cg(283-458) Sc(177-312) | pH173 | this study | Chimera |
| pH438 | Sc(1-176) Cg(283-458) Sc(177-312) | pH173 | this study | Chimera |
| pH440 | Cg(1-44) Sc(43-99) Cg(113-458) Sc(177-312) | pH173 | this study | Chimera |
| pH441 | Sc(1-42) Cg(45-458) Sc(177-312) | pH173 | this study | Chimera |
| pH442 | Sc(1-99) Cg(113-458) Sc(177-312) | pH173 | this study | Chimera |
