## supplementary figures for "Divergence in a Eukaryotic Transcription Factor’s co-TF Dependence Involves Multiple Intrinsically Disordered Regions"

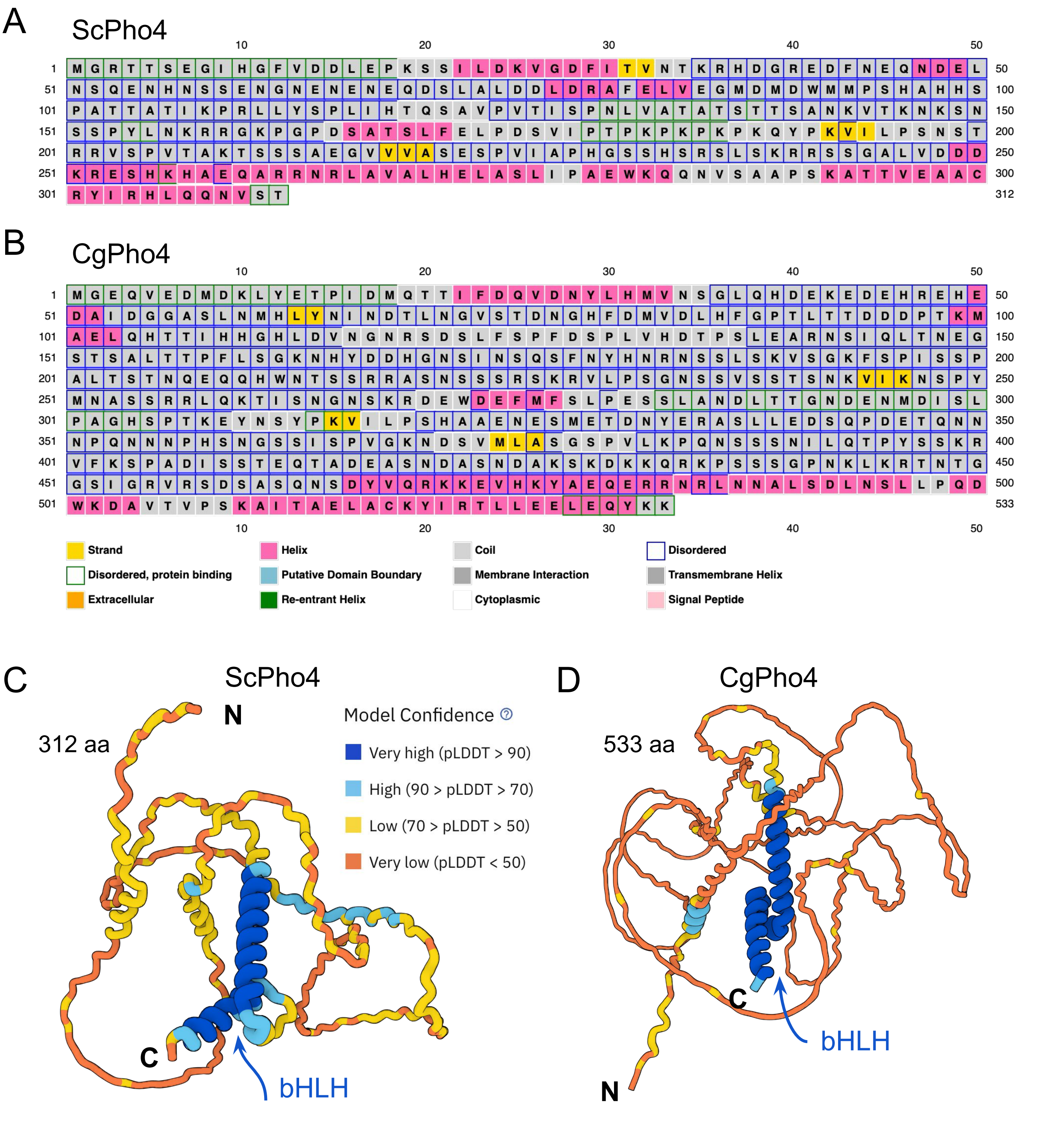


**Supplementary Fig.1 ScPho4 and CgPho4 structural prediction shows both proteins are largely disordered outside the DBD.** (A, B) Secondary structure predictions for ScPho4 and CgPho4 using the PSIPRED 4.0 server. (C, D) AlphaFold2 predicted monomer 3D structure for ScPho4 and CgPho4. Color represents model confidence. It has been shown that low and very low confidence regions overlap significantly and are predictive of protein disorder.

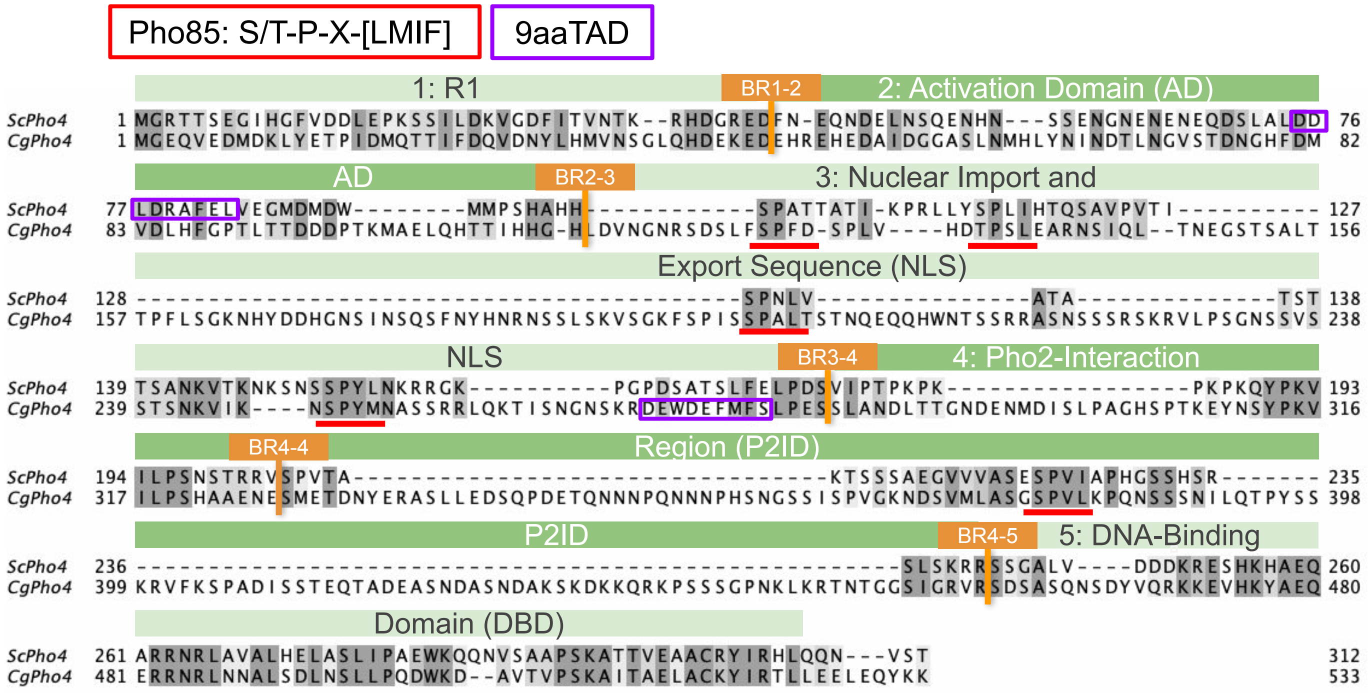


**Supplementary Fig. 2 ScPho4 and CgPho4 alignment and annotation.** Protein sequence alignment was generated using ProbCons with 2 passes of consistency transformation and 100 iterative refinement, and was manually edited to align the known Pho85 recognition motifs (red lines). The shading behind the amino acid symbols represent level of conservation based on BLOSUM62 scores. The functional annotation on the top were based on ScPho4. The orange boxes and texts refer to the breakpoints used for constructing chimeric Pho4 constructs (BR4-4 only used in Fig. 6).

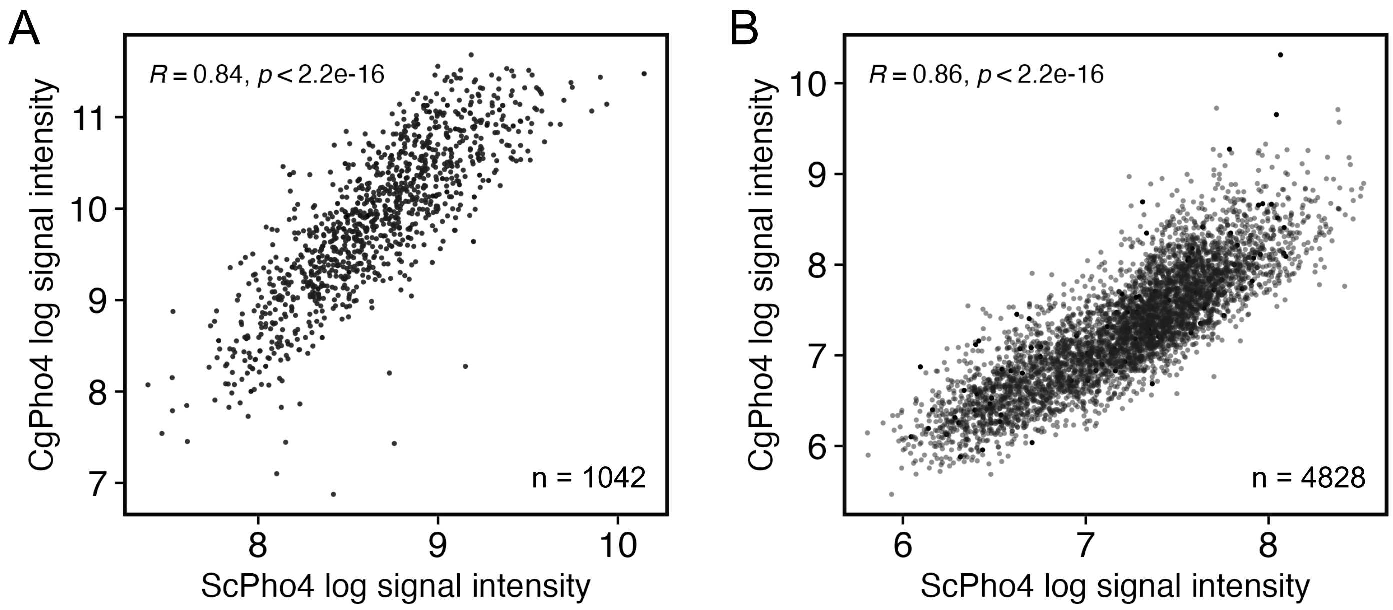


**Supplementary Fig. 3 Genome context Protein Binding Microarray (gcPBM) reveals little divergence in sequence preference between ScPho4 and CgPho4.** (A) 36-bp probes were designed with ChIP-identified binding peaks for both ScPho4 and CgPho4 in their respective genome, centered on the consensus E-box motif, CACGTG, along with the flanking nucleotides. Recombinantly purified full length ScPho4 and CgPho4 were hybridized to separate arrays, and the log signal intensities by the two Pho4 proteins for the same oligo were plotted as a scatterplot. A Spearman's rank correlation coefficient and the *P*-value based on a permutation test for the correlation being 0 (no correlation) are provided at the top of the graph. Number of probes is provided in the lower right. (B) Same as A, except the 36-bp probes were ChIP-identified peaks with a non-consensus E-box motifs (mostly with 1-bp-mismatch to the consensus).

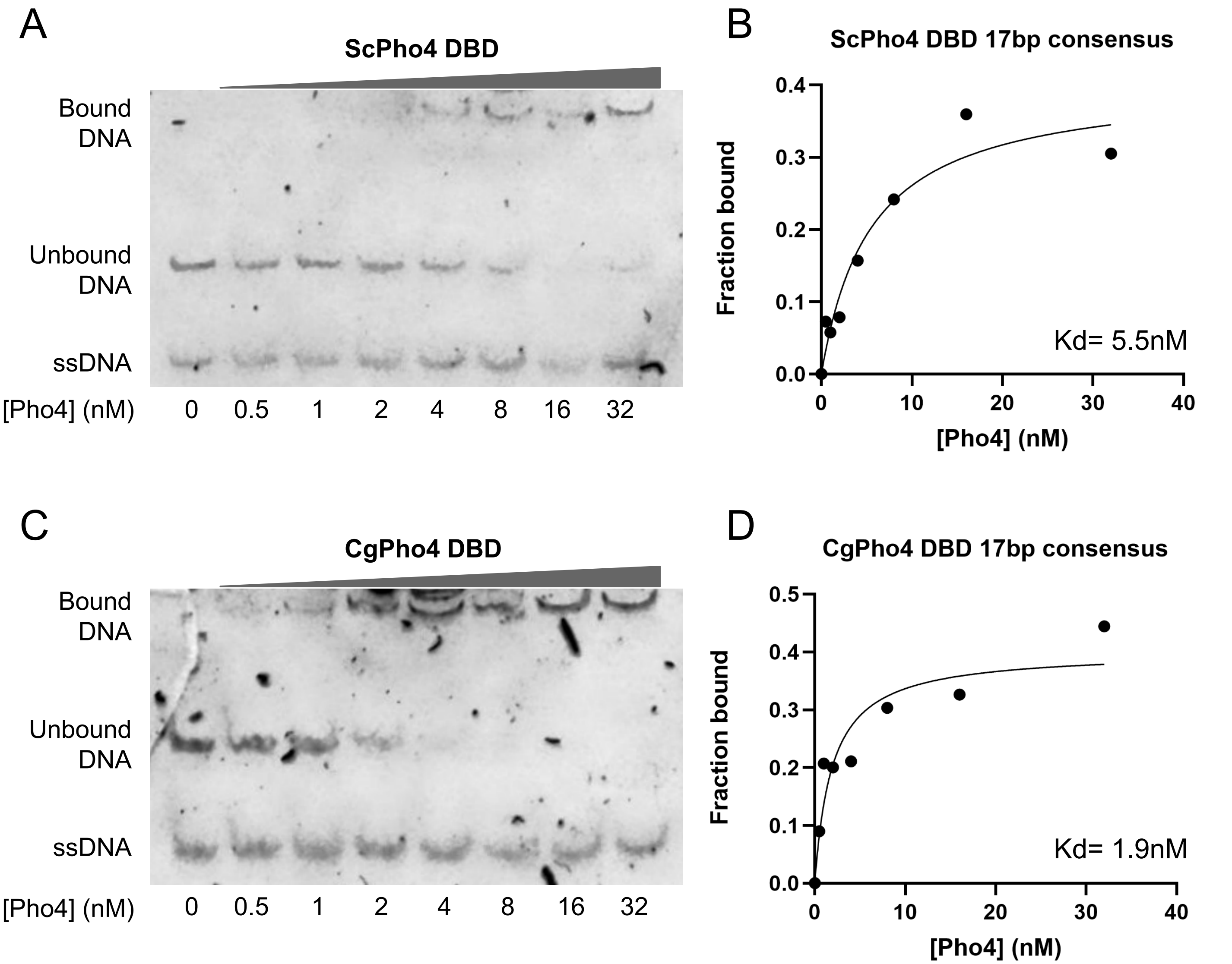


**Supplementary Fig. 4 Electrophoretic Mobility Shift Assay (EMSA) confirms difference in binding affinity to the consensus motif between the two orthologous Pho4 DBDs.** (A) EMSA with ScPho4 DBD and IR700 labeled 17bp consensus DNA. Lanes 2-8 contain 2x diluted protein; lane 1 is a no protein (NP) control. (B) the unbound DNA was quantified from (A) and used calculate the fraction bound normalized to the NP lane. A one-site specific-binding with Hill coefficient model was fit to the data in GraphPad Prism v10. Estimated dissociation constant Kd is shown. (C) and (D) are the same as (A) and (B), but for CgPho4 DBD. Two replicates were performed for each protein and yielded consistent results.

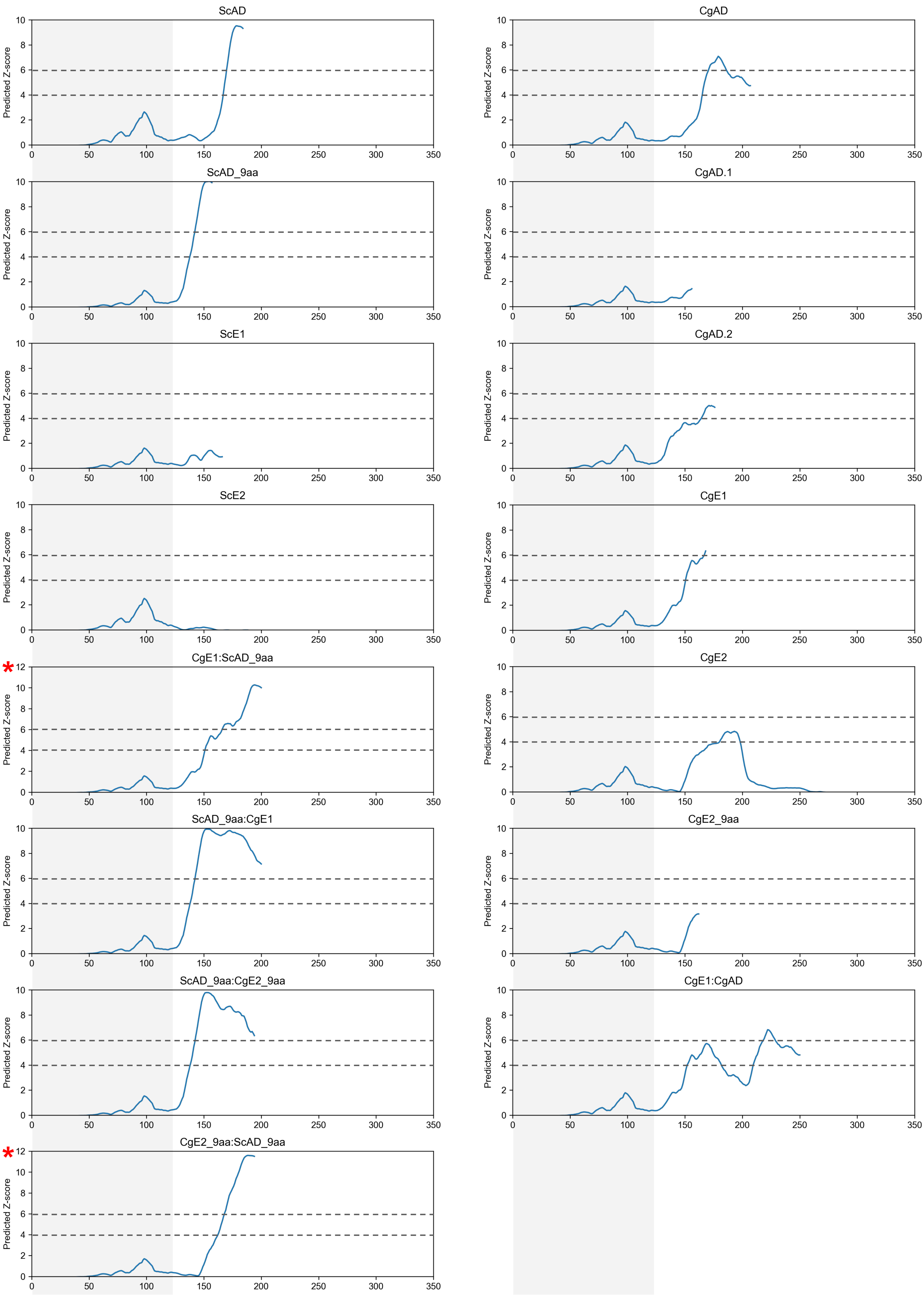
ScAD
CgAD
ScAD_9aa
CgAD.1
ScE1
CgAD.2
ScE2
CgE1
CgE1:ScAD_9aa
CgE2
ScAD_9aa:CgE1
CgE2_9aa
ScAD_9aa:CgE2_9aa
CgE1:CgAD
CgE2_9aa:ScAD_9aa
Predicted Z-score
0 2 4 6 8 10 12
0 50 100 150 200 250 300 350

**Supplementary Fig. 5 PADDLE prediction for synthetic yeast one-hybrid constructs revealed no new regions with activation potential.** PADDLE was run on the full length yeast one-hybrid constructs as was done for the endogenous ScPho4 and CgPho4. Results were plotted using a fixed x axis ranging from 0 to 350 aa even though most constructs were shorter. Like in Fig. 3, the first and last 26 aa had no score due to the design of the prediction algorithm. The greyed region indicates the location of the Gal4 DBD. Its first 22 positions received a Z-score < 0 and hence not visible in the plot. Two constructs had peaks exceeding the Z-score limit of 10. They were plotted with an extended y-axis limit of 0 to 12 (red asterisks). The horizontal dotted lines indicate the thresholds for medium and strong activation potentials, respectively.

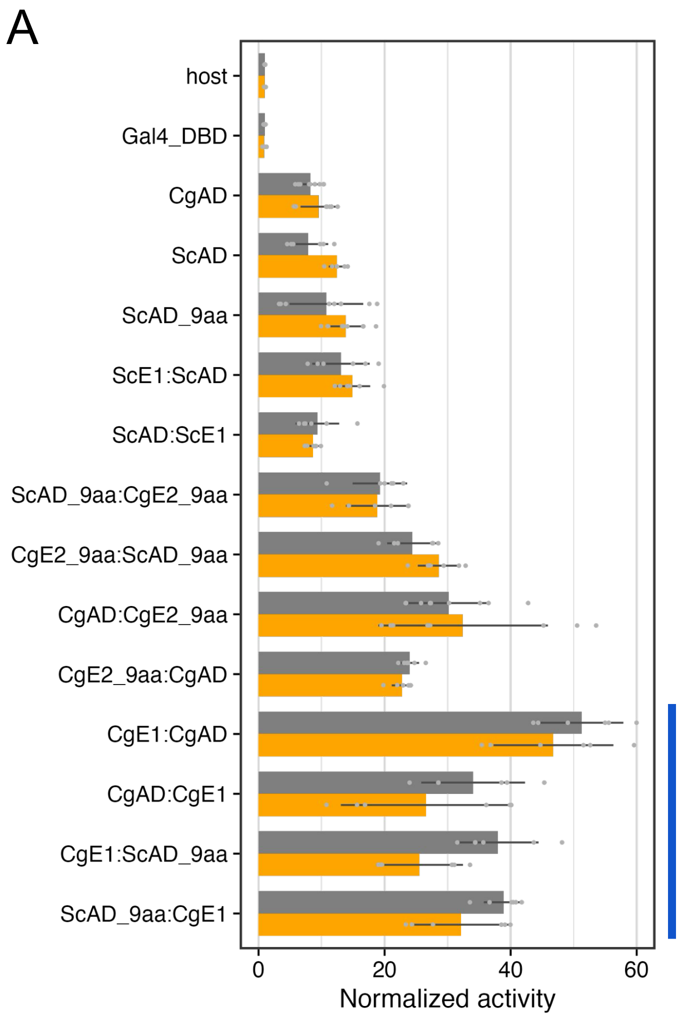


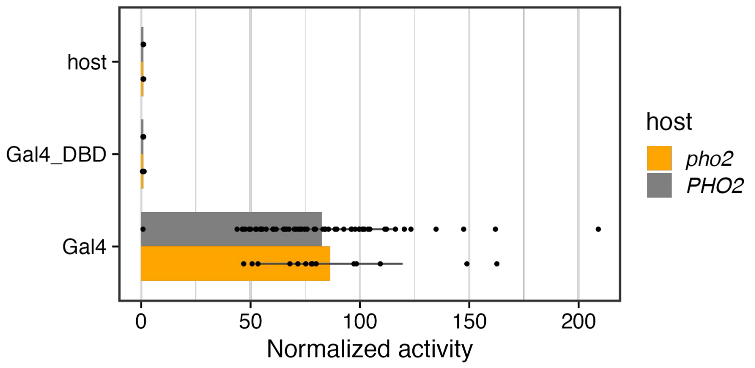


B

**CgE1 boost effect without *pho2***

| Group | ∆mean | raw P |
|---|---|---|
| CgE1:CgAD vs CgAD | 37.3 | 3x10$^{75}$ |
| CgAD:CgE1 vs CgAD | 17.0 | 0.002 |
| CgE1:ScAD_9aa vs ScAD_9aa | 11.7 | 0.001 |
| ScAD_9aa:CgE1 vs ScAD_9aa | 18.3 | 1x10$^{-5}$ |

**Supplementary Fig. 6 Testing Pho2-dependence for the activation potential in the Yeast-1-Hybrid (Y1H) assay.** (A) Y1H was performed both with and without *PHO2* (gray and orange bars, respectively). “Host” = no plasmids All names are as specified in Fig. 3 of the main text. Bars represent the means of n>3 replicates; individual data points are shown as dots; lines are the standard deviations. Student’s t-tests were used to compare the activity of each construct with vs without Pho2. None of the tested constructs was significant at a 0.05 level after Bonferroni-Holm correction. Full length Gal4 was tested in the same experiment and plotted on the right to avoid compressing the scale. (B) Constructs highlighted by the blue vertical line in the left panel contain CgE1 and were tested for the boosting effect without Pho2 by comparing their activity against the corresponding AD alone. Two-sided student’s t-tests were performed and the difference in the mean and raw *P*-values were reported.

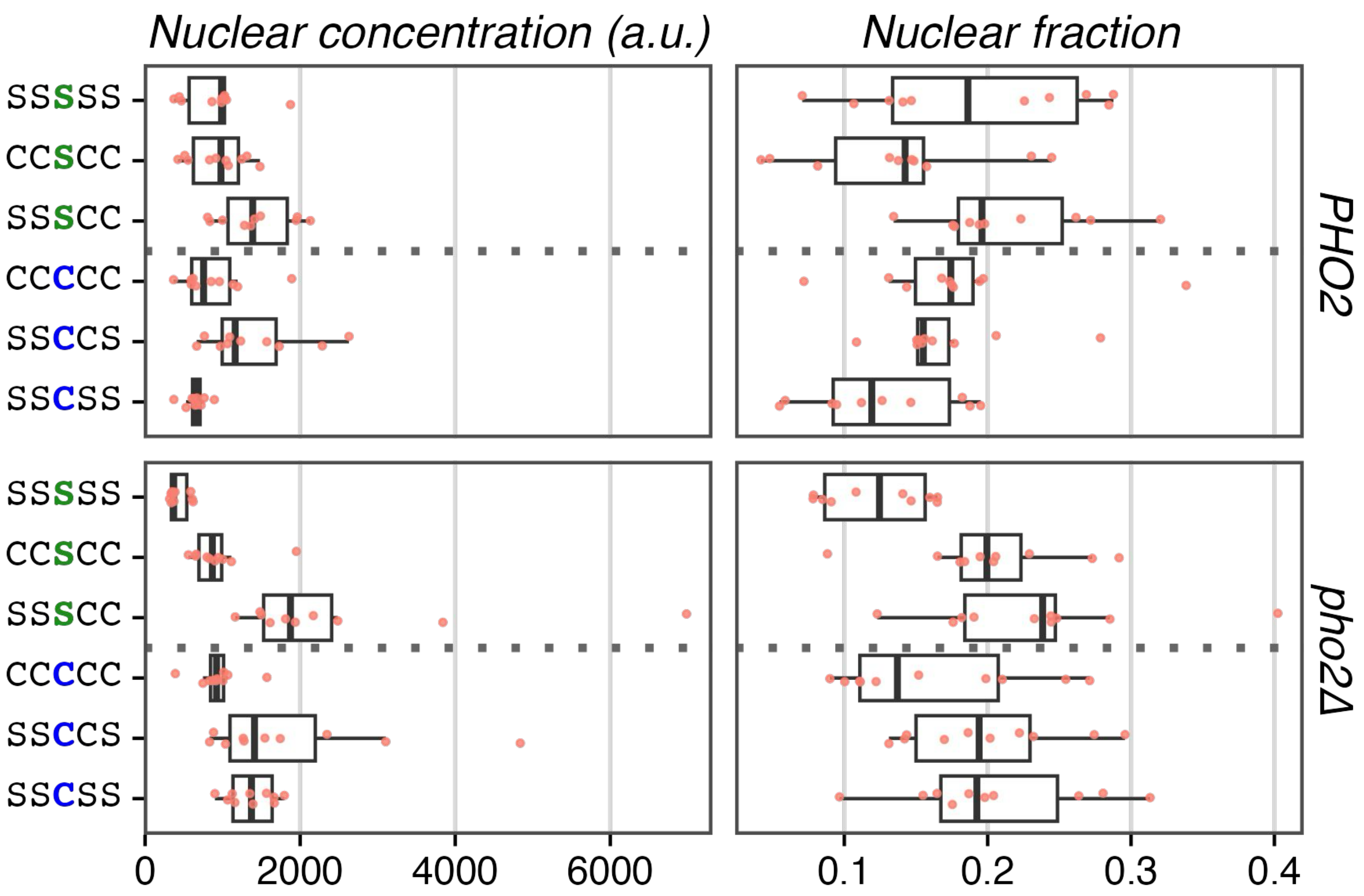


**Supplementary Fig. 7 NLS from ScPho4 and CgPho4 conferred similar levels of nuclear localization.** Six Pho4 constructs with either NLS:Sc or NLS:Cg were quantified for nuclear Pho4 concentration and fraction of Pho4 in the nucleus, in either *PHO2* or *pho2*Δ host background. Endogenous or chimeric Pho4 protein levels were quantified by the C-terminal mNeon tag. Nucleus was labeled by DAPI. Nuclear concentration was estimated using the average gray intensity inside the nucleus. Nuclear fraction was calculated by dividing the Integrated density inside the nucleus by the total integrated density in the whole cell. A total of 10 cells were randomly selected from the bright field image per construct in each background. Individual estimates are shown as dots and a boxplot, where the box represents the interquartile range, the middle line represents the median and the whiskers 1.5 times the interquartile range. ANOVA test found no significant differences for either the identity of the NLS ("Cg" or "Sc") or Host (*PHO2* or *pho2*Δ) factors (F-test raw *P* = 0.41 and 0.15, respectively).

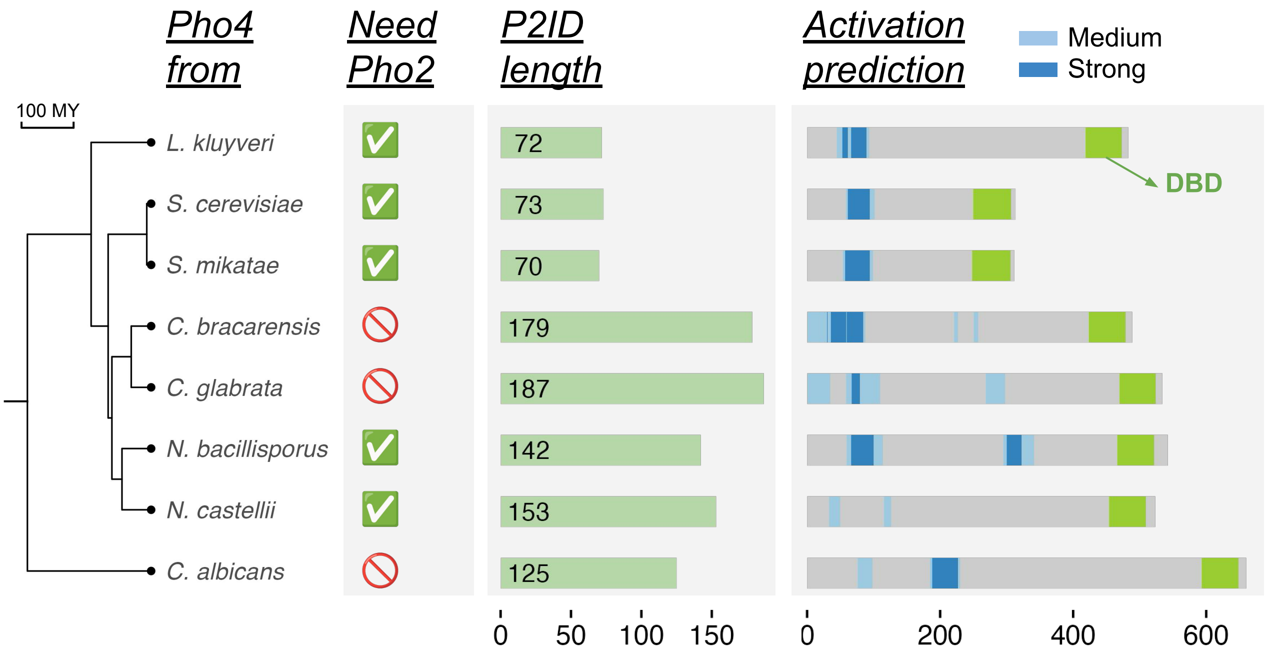


**Supplementary Fig. 8 Evolutionary patterns of AED and P2ID length among Pho4 orthologs.** Time-calibrated species tree on the left is based on Shen et al. 2018 (PMID: 30415838). Pho4 is a single gene family in all budding yeasts, including the 8 species shown. Among these Pho4 orthologs, three showed reduced dependence on Pho2 based on He et al. 2017 (PMID: 28485712). Middle: P2ID length is calculated based on the multiple sequence alignment constructed using ProbCons. Right: activation potential is predicted using PADDLE. Medium (Z-score between 4 and 6) and strong (Z-score > 6) activation potential regions are shown as light and dark blue stripes. The bHLH DBD is shown as green box for reference.

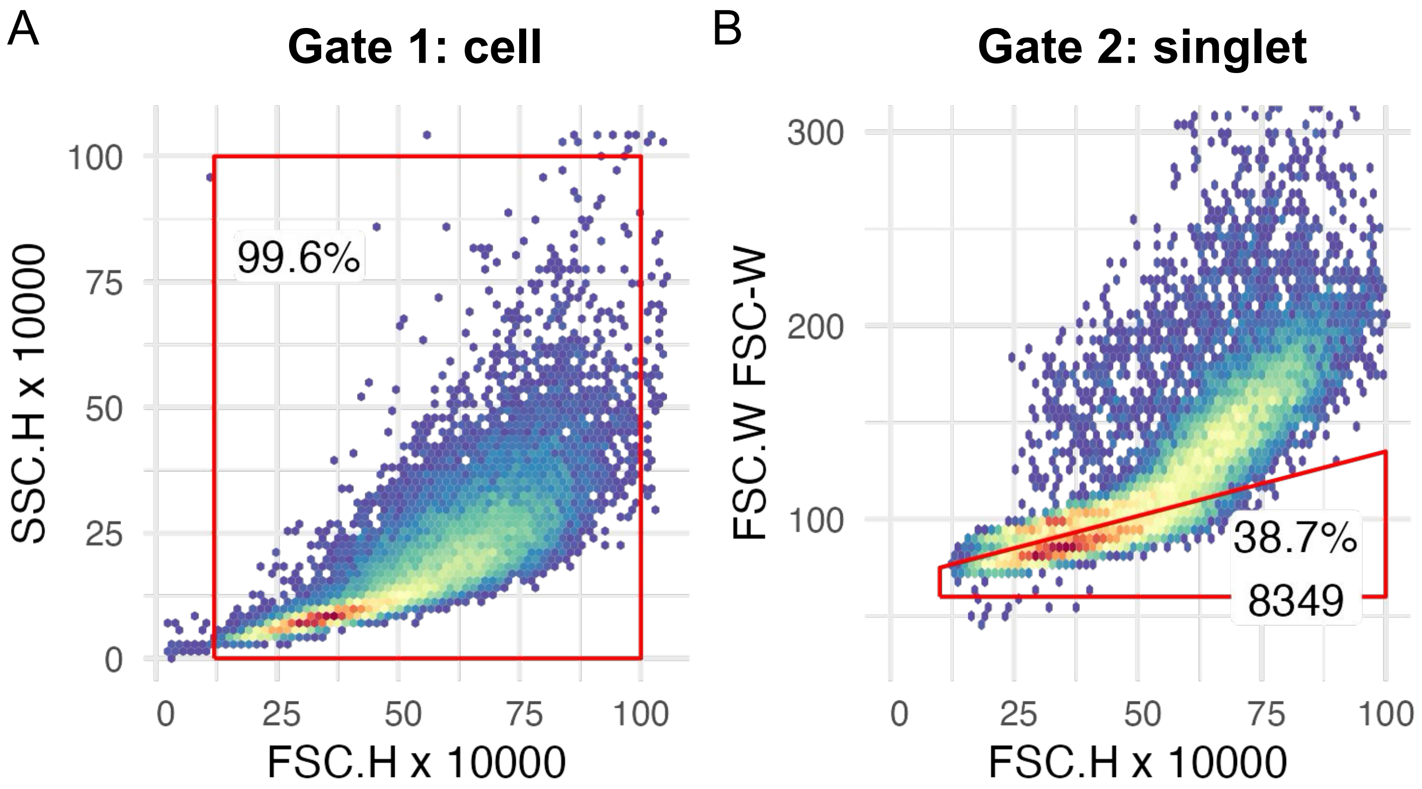


**Supplementary Fig. 9 Example gating strategy for flow cytometry data.** (A) All events were plotted on FSC.H and SSC.H. A rectangular gate is used to exclude non-cell events. (B) Events within the first gate ("cell") were plotted on FSC.H and FSC.W. Singlets (single cell event as opposed to doublets or multilets) were selected by excluding events with a higher FSC.W given the same FSC.H.
